# Movement abilities and brain development in preschoolers born very preterm

**DOI:** 10.1101/734319

**Authors:** Holly M. Hasler, Martha G. Fuller, Yvonne E. Vaucher, Timothy T. Brown, Joan Stiles, Anders M. Dale, Terry L. Jernigan, Natacha Akshoomoff

## Abstract

**Aim:** To examine how healthy preschoolers born very preterm (VPT) with and without significant movement impairments differ from full term (FT) controls in subcortical brain volume measures and white matter diffusion properties.

**Method:** A case-control, observational study of fifty-four VPT-born and 32 FT-born children were administered the Movement Assessment Battery for Children – Second Edition (MABC-2) and underwent MRI within 6-months of starting kindergarten. Selected subcortical structural volumes, fractional anisotropy (FA), and mean diffusivity (MD) of selected white matter tracts were compared across VPT children with movement impairments (VPT-abnormal), and VPT and FT children without movement impairments.

**Results:** The VPT-abnormal group had higher MD in the corpus callosum and inferior frontal-occipital fasciculus and lower FA in the anterior thalamic radiations, corpus callosum, and cingulum than the FT group. The forceps major was particularly affected in the VPT-abnormal group compared with the VPT and FT groups without movement impairments. Both VPT groups had reduced brainstem and cerebellar white matter volumes and larger lateral ventricles compared to the FT group.

**Interpretation:** Movement impairments in healthy VPT preschoolers were associated with more abnormalities in white matter integrity and reduced subcortical brain volumes most likely reflecting a greater extent of white matter damage associated with their very preterm birth.

## Movement abilities and brain development in preschoolers born very preterm

Children born very preterm (VPT), at gestational age less than 33 weeks, are at increased risk for health problems, behavioral difficulties, and cognitive impairments. Motor deficits are prominent with some studies reporting almost 40% of extremely preterm (< 28 weeks) children displaying mild motor deficits and up to 10% developing cerebral palsy (Adams-Chapman et al., 2018; de Kieviet, Piek, Aarnoudse-Moens, & Oosterlaan, 2009; Edwards et al., 2011; Spittle, Cameron, Doyle, Cheong, & Victorian Infant Collaborative Study, 2018; Williams, Lee, & Anderson, 2010).

Preterm birth increases the risk for periventricular white matter injury in the neonatal period which can contribute to more diffuse alterations in white matter pathways and distal gray matter (Volpe, 2009). Compared to their full term (FT) born peers, reduced volumes of the thalamus, cerebellar white matter, and anterior cingulate were reported in one study of toddlers born VPT (Lowe et al., 2011). In a number of studies of older VPT children and adolescents, reduced volumes of the corpus callosum, basal ganglia, thalamus, brainstem, ventral diencephalon, and cortical white matter regions volumes are commonly found (J. F. de Kieviet et al., 2014; de Kieviet, Zoetebier, van Elburg, Vermeulen, & Oosterlaan, 2012; Lax et al., 2013; Solsnes et al., 2016).

Injury to the regions that are typically most affected by preterm birth may help explain the prevalence of motor dysfunction in children born VPT. White matter integrity and injury severity in infants scanned near-term equivalent age have been shown to be related to motor abilities assessed later in development (Rose et al., 2015; Schadl et al., 2018; Spittle et al., 2011). In toddlers, fractional anisotropy (FA) of the corpus callosum, cingulum, fornix, and uncinate fasciculus were correlated with movement skills (Counsell et al., 2008). Movement abilities in older children and adolescents have been shown to be correlated with the volume of the corpus callosum, cerebellar white matter, and cerebellar gray matter (de Kieviet et al., 2012). In a study of white matter integrity at 8-years of age, VPT children who scored <15^th^ percentile on the Movement Assessment Battery for Children (MABC) showed FA differences in widespread white matter tracts including in the cingulum, corticospinal tract (CST), forceps major, and forceps minor (Jorrit F. de Kieviet et al., 2014).

Motor skill assessments at age 4 have been shown to be more reliable than assessments conducted at age 2, which tend to underestimate movement difficulties in VPT children without cerebral palsy (Griffiths et al., 2017; Spittle et al., 2013). Assessment of motor skills just prior to entering kindergarten may identify persistent motor difficulties that may be associated with cognitive and academic challenges (Brostrom, Vollmer, Bolk, Eklof, & Aden, 2018; Maggi, Magalhaes, Campos, & Bouzada, 2014).

The objective of this study was to examine the movement skills and differences between subcortical volumes and white matter fiber volume and diffusion properties between VPT-born preschoolers with impaired movement assessments and VPT and FT preschoolers with movement assessments in the normal range. To date no published studies have examined how movement abilities relate to brain structure when both are measured at preschool age in VPT children without major neurodevelopmental disorders or other significant functional impairments.

## Method

### Participants

Participants were healthy children born VPT (*n* = 54) and FT (*n* = 32) who were part of an ongoing, longitudinal study of cognitive and academic skills (Hasler & Akshoomoff, 2017). Demographic characteristics are shown in Table 1. A convenience sample of VPT children was recruited primarily from the neurodevelopmental follow-up program of a tertiary NICU (UC San Diego). Exclusion criteria included history of severe neonatal brain injury (i.e., Grade 3-4 intraventricular hemorrhage, cystic periventricular leukomalacia, moderate-severe ventricular dilation), known genetic abnormalities likely to affect development, and/or acquired neurological disorder unrelated to preterm birth. Children born FT were recruited from the UC San Diego Center for Human Development database of parents who consented to be contacted for research studies and had no history of neurological, psychiatric, or developmental disorders. VPT and FT born children were primarily English speaking with a WPPSI Full Scale IQ > 75, without significant auditory or visual deficits, or contraindications to MRI. One child born FT was excluded due to a brain abnormality identified after review by our study neuroradiologist. Three children born VPT and two children born FT were excluded who had excessive movement artifacts in their diffusion-weighted images or errors in MRI segmentation. Six children born FT were excluded for MABC-2 scores in the impaired range.

**Table 1.**
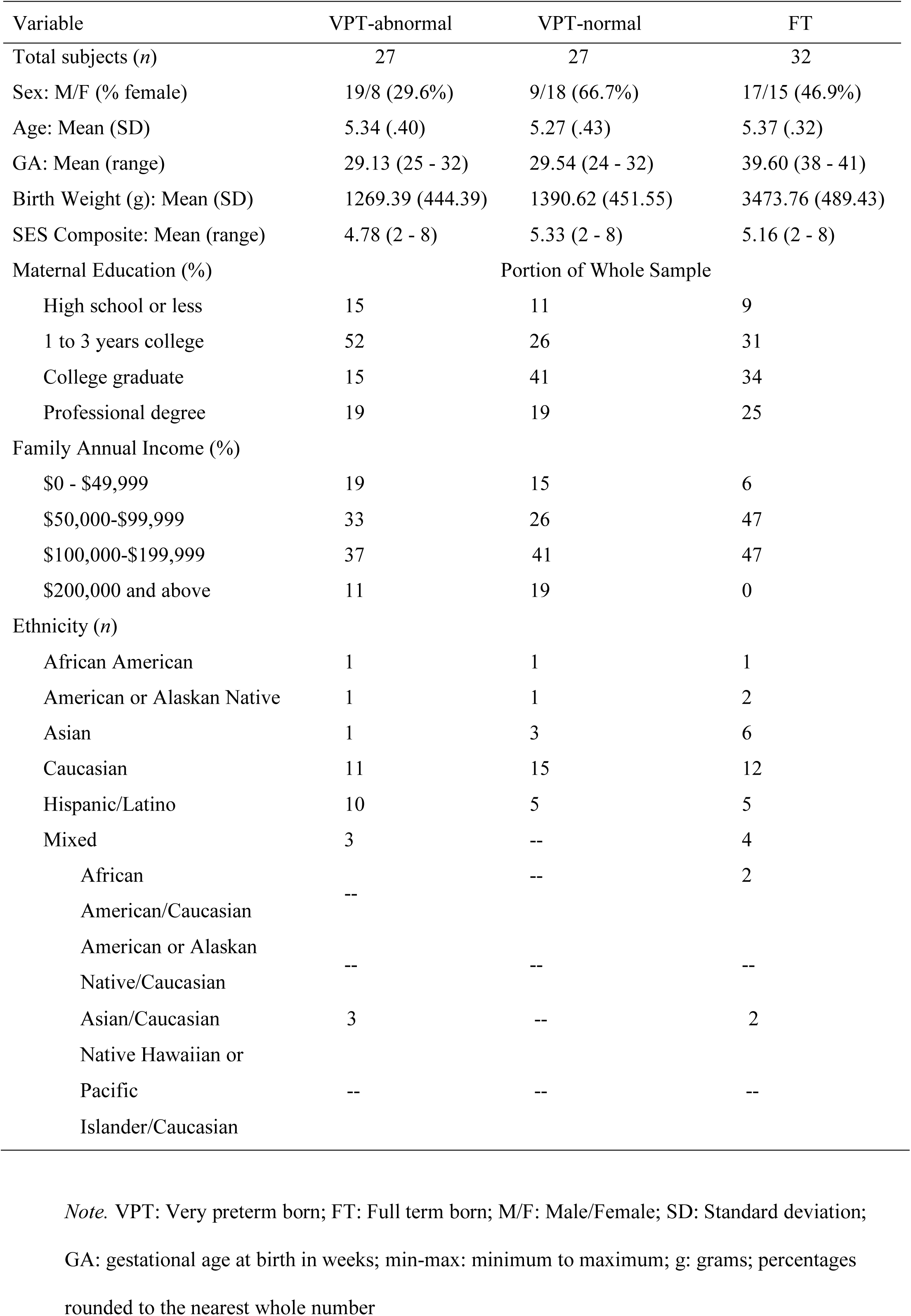
Participant Demographics

The Institutional Review Board at UC San Diego approved all procedures, and each participant’s legal guardian gave written informed consent.

### Movement Assessment Battery for Children – Second Edition (MABC-2)

The MABC-2 (Henderson, Sugden, & Barnett, 2007), a norm-referenced assessment that measures manual dexterity, balance, and visual-motor coordination through a series of motor tasks, was administered to all children by a single examiner (MGF). The three MABC-2 subscale scores (Aiming and Catching, Balance, and Manual Dexterity) are combined to form a Total Score. Scaled scores for the individual subscales and the Total Score range from 1 to 19, with higher scores representing better performance. It is recommended that children with Total Scores at or below the 15^th^ percentile receive ongoing monitoring of their movement skills. For this study, impairment was defined by a Total Score of ≤ 6, equivalent to performance below the 15^th^ percentile for age. Subscale scores of ≤ 6 are at or below the 9^th^ percentile for age.

### Wechsler Preschool and Primary Scale of Intelligence – Fourth Edition (WPPSI – IV)

The WPPSI-IV is a broad measure of cognitive abilities used in children and includes indices of verbal comprehension, visual spatial abilities, fluid reasoning, working memory, and processing speed (Wechsler, 2012). The WPPSI-IV was administered by trained, reliable examiners to assess overall cognitive functioning for inclusion in the study.

### Brain Imaging

Each child underwent a mock MRI to familiarize them with the procedure, with the study MRI on a separate day.

Data were collected on a General Electric Discovery MR750 3.0 Tesla scanner with an 8-channel phased-array head coil at the Center for Functional MRI at UC San Diego. The imaging protocol included 1) a three-plane localizer; 2) a 3D T1-weighted inversion prepared RF-spoiled gradient echo scan using prospective motion correction (PROMO) for cortical and subcortical segmentation; 3) a 3D T2-weighted variable flip angle fast spin echo scan for detection and quantification of white matter lesions and segmentation of CSF; 4) a high angular resolution diffusion imaging (HARDI) scan with integrated B_0_ distortion correction (DISCO), for segmentation of white matter tracts and calculation of diffusion parameters. All data were inspected for quality during collection and at all stages of processing.

Data were processed at the UC San Diego Center for Multimodal Imaging and Genetics (CMIG) using gradient nonlinearity correction, co-registration of images, and EPI B_0_ distortion correction. In addition, images were processed using and automated probabilistic, atlas-based analysis of white matter tracts (Hagler et al., 2009) and cortical and subcortical segmentation was performed using FreeSurfer version 5.3.0 (Fischl et al., 2002). The imaging protocol was developed specifically for data collection with children as part of the Pediatric Imaging Neurocognition and Genetics project (PING) (Jernigan et al., 2016).

### Statistical Analyses

All statistical analyses were completed in SPSS, version 23. (SPSS, 2014) Children were categorized into three groups by their Total MABC-2 score: “VPT-abnormal” with scores in the impaired range (scaled score ≤ 6 and <15^th^ percentile); “VPT-normal” and FT children with scores in the normal range (scaled scores ≥ 7 and > 15^th^ percentile).

Demographic variables were compared between groups using Pearson χ^2^ and *t*-tests. A single SES value was calculated as a combination of parent-reported household income (4 levels) and years of maternal education (4 levels). This resulted in a value from 1 to 8 for each child.

Group whole brain and ventricle volumes were inspected for outliers using boxplots; none were found. MANCOVAs were used to compare values derived from brain measures between groups. Brain structure and white matter tract volumes were compared controlling for sex, SES, and FreeSurfer-derived intracranial volume (ICV). Diffusion measure comparisons (FA and MD) controlled for sex and SES. When evaluating the assumptions of the MANCOVA, Box’s M was significant for both models. Therefore, Pillai’s Trace (*V*), was used as a more robust estimation of overall effect. Individual neuroanatomical regions were entered simultaneously into the model. Bonferroni corrected, *a priori* pair-wise t-tests were conducted to elucidate differences among groups. There were no *a priori* hypotheses regarding left versus right hemisphere differences, therefore values were summed across hemispheres for volumes and averaged for DTI values to reduce the number of multiple comparisons. However, some previous studies have reported hemispheric differences; thus, follow-up group comparisons of each hemisphere separately were conducted using t-tests with Bonferroni correction.

## Results

The three groups did not differ in age (*F* (2,81) = .681, *p=*.51) or SES (χ^2^ = 9.90, *p* = .63). The VPT-abnormal and VPT-normal groups did not differ in gestational age (*t* (52) = -.647, *p* = .52) or birthweight (*t* (52) = -.678, *p* = .51). The total VPT group (VPT-abnormal + VPT-normal) was 48.1% female, which is similar to the FT group (χ^2^ = .013, *p* = .909, 46.9% female). However, the VPT-abnormal and VPT-normal groups differed significantly in proportion of females (χ^2^ = 7.42, *p* = .006). Subsequent analyses were conducted with sex as a covariate in order to better account for possible sex differences. FSIQ was not a significant factor in the omnibus MANCOVA for brain volumes (*V* = .120, *F* (34, 124) = .787, p *=* .662, ηp^2^ = .120), or the diffusion model (*V* = .174, *F* (17, 65) = .804, p *=* .682, ηp^2^ = .174), including all pair-wise comparisons. In addition, FSIQ was not a significant between-subject predictor in either model. Therefore, FSIQ was not included in the final model as a covariate.

Results of the one-way ANOVA demonstrated Total MABC-2 scores as well as the three subtest scores were significantly different between the three groups (Table 2). In the post-hoc pairwise analyses, after Bonferroni correction, each subtest score was significantly lower in the VPT-abnormal group compared to VPT-normal (all *p*s < .001) and FT groups (all *p*s < .001). Additionally, the VPT-normal group performed significantly worse on the Balance subtest (*p =* .01) than the FT group. The VPT-abnormal group had significantly lower Full Scale IQ scores (FSIQ) than the FT group, although all three group mean scores were within the average range (see Table 2). Total MABC-2 scaled score was significantly correlated with FSIQ across all the children in the study (*r =* .252, *p =* .019).

**Table 2.** Performance on movement measure and FSIQ by group

| | VPT-abnormal | VPT-normal | FT | ANOVA | $\eta^2$ |
| --- | --- | --- | --- | --- | --- |
|  | M (SD) | M (SD) | M (SD) | *F* (*p*) |  |
| **MABC-2** |  |  |  |  |  |
| Aim & Catch | 6.70 (2.74)<br>a, b | 9.96 (2.19) | 10.31<br>(3.02) | 15.28 (<.001) | .269 |
| Balance | 5.48 (1.81)<br>a, b | 8.74 (2.86)<br>a | 10.71<br>(3.24) | 27.00 (<.001) | .394 |
| Manual Dexterity | 4.74 (2.14)<br>a, b | 9.67 (2.42) | 9.63 (3.20) | 31.40 (<.001) | .431 |
| Total | 4.26 (1.51)<br>a, b | 9.11 (1.74) | 10.13<br>(2.93) | 57.16 (<.001) | .579 |
| **WPPSI-IV** |  |  |  |  |  |
| FSIQ | 99.48<br>(12.83)<sup>a</sup> | 103.52<br>(12.76) | 108.31<br>(9.77) | 4.17 (.019) | .091 |

*Note.* Pairwise comparisons: <sup>a</sup>: Significantly different from FT Group; <sup>b</sup>: significantly different from VPT-normal Group; VPT: Very preterm born; FT: Full term born; M: Mean; SD: Standard Deviation; MABC-2: Movement Assessment Battery for Children – Second Edition; Aim & Catch: Aiming and Catching; WPPSI-IV: Wechsler Preschool and Primary Scale of Intelligence - Fourth Edition; FSIQ: Full Scale IQ. Two FT-born children were administered the WPPSI-III, therefore their scores are not included

When evaluating the assumptions of the MANCOVA, Box’s M was significant for both models. Therefore, Pillai’s Trace (*V*), was used as a more robust estimation of overall effect. There was a statistically significant OMNIBUS difference across the brain volume measures (*V* = .614, *F* (24, 140) = 2.58, p < .001, ηp^2^ = .307). Post-hoc analyses indicated group differences for volumes of the brainstem (*F*(2, 84) = 10.25, *p* < .001, ηp^2^ = .204), lateral ventricles (*F*(2, 84) = 7.17, *p =* .001, ηp^2^ = .152), forceps major (*F*(2, 84) = 6.48, *p =* .002, ηp^2^ = .139), thalamus (*F*(2, 84) = 4.44, *p =* .01, ηp^2^ = .100), and cerebellar white matter (*F*(2, 84) = 5.78, *p =* .005, ηp^2^ = .126). Follow-up testing between groups is presented in Table 3. The FT group had a higher brainstem, lateral ventricle, and cerebellum volume than both VPT groups. FT had higher volume than only the VPT-normal group. Both the VPT-normal and FT groups had higher forceps major volume than the VPT-abnormal group. Volume of the other white matter regions entered in the model (corpus callosum, forceps minor, anterior thalamic radiations (ATR), and cortical-spinal tract), and subcortical structures (ventral diencephalon and basal ganglia) were not significant predictors of group.

**Table 3.** Regional brain volume and diffusion measures by group

| Region | VPT-abnormal | VPT-normal | FT |
| --- | --- | --- | --- |
| Vol Brainstem | 16026.27 (265.25)<sup>a</sup> | 16131.93 (265.76)<sup>a</sup> | 17430.02 (236.58) |
| Vol Lateral Ventricles | 14145.22 (1572.12)<sup>a</sup> | 13044.51 (1575.14)<sup>a</sup> | 6943.41 (1402.18) |
| Vol Thalamus | 13353.89 (161.70) | 13228.88 (162.01)<sup>a</sup> | 13826.30 (144.22) |
| Vol Cerebellum WM | 21230.03 (516.92)<sup>a</sup> | 21566.21 (517.92)<sup>a</sup> | 23361.36 (461.05) |
| Vol Forceps Major | 10045.89 (305.54)<sup>a, b</sup> | 11299.65 (306.13) | 11442.70 (272.51) |
| MD Forceps Major | .977 (.009)<sup>a, b</sup> | .934 (.009) | .923 (.008) |
| MD Corpus Callosum | .913 (.008)<sup>a</sup> | .895 (.008) | .881 (.007) |
| FA Forceps Major | .492 (.008)<sup>a, b</sup> | .520 (.008) | .533 (.007) |
| FA ATR | .338 (.004)<sup>a</sup> | .347 (.004) | .353 (.004) |
| FA Cingulum | .375 (.006)<sup>a</sup> | .392 (.006) | .406 (.005) |

*Note.* Marginal means calculated from the MANCOVA controlling for SES, sex, and ICV for volumes and SES and sex for MD and FA; Volume measures presented as adjusted mean mm<sup>3</sup> after controlling for total intracranial volume (standard error); Post-hoc comparisons using Bonferroni correction for multiple comparisons: <sup>a</sup>significantly different from FT group; <sup>b</sup>significantly different from VPT-normal group; ICV: intracranial volume; Vol: volume; WM: white matter; MD: mean diffusivity; FA: fractional anisotropy;

The omnibus MANCOVA revealed a significant group difference across the diffusion measures (*V* = .617, *F* (34, 132) = 1.73, *p =* .01, ηp^2^ = .309). Post-hoc analyses indicated group differences for FA (*F*(2, 84) = 6.85, *p =* .002, ηp^2^ = .145) and mean diffusivity (MD) ((2, 84) = 9.99, *p* < .001, ηp^2^ = .198) of the forceps major, MD of the corpus callosum (*F*(2, 84) = 4.27, *p =* .017, ηp^2^ = .095), FA of the ATR (*F*(2, 84) = 3.89, *p =* .024, ηp^2^ = .088), and FA of the cingulate (*F*(2, 84) = 7.35, *p =* .001, ηp^2^ = .154). Follow-up testing between groups is presented in Table 3. VPT-abnormal group had significantly higher MD and lower FA in the forceps major than the VPT-normal and FT groups. The VPT-normal group had significantly higher MD and lower FA in the corpus callosum, ATR, and cingulum, compared to the FT group. Diffusion measures of the inferior frontal-occipital fasciculus, thalamus, basal ganglia, superior cortical striatal fibers, and CST were not significant predictors of group.

Planned, follow-up group comparisons of separate hemispheres were conducted. Regions that were found to have significant associations with group in follow-up testing are presented in Table 4. Compared with the FT group, the volume of the left thalamus was significantly smaller in the VPT-abnormal (*p =* .021) and VPT-normal (*p* = .027) groups and the right thalamic volume (*p =* .007) was significantly smaller in the VPT-normal group. The remaining measures were significant for both left and right hemispheres. Cerebellar white matter volume was significantly smaller in both the left (*p =* .003) and right (*p =* .005) hemisphere for the VPT-abnormal group, as well as in the left (*p =* .027) and right (*p =* .007) hemisphere for the VPT-normal group. With regard to the diffusion measures, both left (*p =* .02) and right (*p =* .04) FA of the ATR as well as left (*p =* .001) and right (*p =* .02) cingulum FA were significantly reduced in the VPT-abnormal group relative to both the VPT-normal and FT groups.

**Table 4.** Estimated marginal means for follow-up separate hemisphere analyses of significant pairwise group comparisons

| Region | VPT-abnormal | VPT-normal | FT |
| --- | --- | --- | --- |
| Vol R Thalamus | 6739.65 (92.19) | 6598.95 (92.36)<sup>a</sup> | 6944.45 (82.22) |
| Vol L Thalamus | 6614.24 (84.97)<sup>a</sup> | 6629.93 (85.14)<sup>a</sup> | 6881.86 (75.79) |
| Vol R Cerebellum WM | 10693.16 (267.08)<sup>a</sup> | 10738.68 (267.60)<sup>a</sup> | 11732.21 (238.21) |
| Vol L Cerebellum WM | 10536.87 (265.21)<sup>a</sup> | 10827.53 (265.72)<sup>a</sup> | 11629.15 (236.54) |
| FA R ATR | .331 (.004)<sup>a</sup> | .339 (.004) | .345 (.004) |
| FA L ATR | .344 (.005)<sup>a</sup> | .354 (.005) | .361 (.004) |
| FA R Cingulum | .363 (.006)<sup>a</sup> | .371 (.006) | .385 (.006) |
| FA L Cingulum | .387 (.008)<sup>a</sup> | .412 (.008) | .426 (.007) |

*Note.* Marginal means calculated from the MANCOVA controlling for SES, sex, and ICV for volumes and SES and sex for MD and FA; Volume values presented as adjusted mean mm<sup>3</sup> after controlling for intracranial volume (standard error); Post-hoc comparisons using Bonferroni correction for multiple comparisons: <sup>a</sup>: significantly different from FT group; <sup>b</sup>: significantly different from VPT-normal group; ICV: intracranial volume; Vol: volume; WM: white matter; MD: mean diffusivity; FA: fractional anisotropy.

## Discussion

Half of this sample of healthy, preschool-aged children born VPT had significant deficits on a standardized test of movement abilities. The VPT-abnormal children in this study demonstrated deficits in movement abilities across all subsections of the MABC-2 including gross motor (Balance), visual-motor coordination (Aiming & Catching), and fine motor dexterity skills (Manual Dexterity). Despite having normal MABC-2 total scores, the children in the VPT-normal group had significantly lower scores on the Balance subscale compared with the FT group, a concerning finding that requires additional evaluation. The higher proportion of males in the VPT-abnormal group adds further evidence that boys born preterm tend to be at higher risk for motor difficulties than girls (Arnaud, Daubisse-Marliac, White-Koning, & et al., 2007; Kuban et al., 2016).

Previous studies have demonstrated a difference in birth characteristics between children born VPT with and without movement difficulties (de Kieviet et al., 2009), though these studies often included children with more risk factors than in this study. Our cohort of VPT children were intentionally selected as “healthy” based on a relatively benign neonatal course. There were no differences in gestational age, birth weight, or NICU course between the VPT-abnormal and VPT-normal groups. The VPT children had no significant neurologic or medical complications related to prematurity and no significant cognitive deficits.

Abnormal motor skills were associated with decreased subcortical volumes and abnormal diffusion measures (higher MD and lower FA) across many white matter tracts. Significant brain region volumes and white matter diffusion property differences were found bilaterally between VPT-born preschoolers with motor impairments, compared to VPT without motor impairments and their FT-born peers. The areas related to motor problems are those that are most vulnerable to the effects of preterm birth (Volpe, 2009) and included subcortical structures, corpus callosum, anterior thalamic radiations, and cingulum fibers.

The abnormalities in white matter microstructure and subcortical brain volumes found in the VPT-abnormal group (i.e., forceps major, cingulum, cerebellum) have been demonstrated in older children with motor impairments who were born VPT (Jorrit F. de Kieviet et al., 2014; de Kieviet et al., 2012). We demonstrated many of the same findings here, though de Kieviet et al. (Jorrit F. de Kieviet et al., 2014) found a more extensive network of association tracts were related to motor impairments in 7-to 8-year-old children born VPT. The younger children in the current study have not yet reached the peak of the myelination process for association fibers (Brown et al., 2012) which may explain why some group differences were not found in our young sample.

In our study the white matter properties of the corpus callosum, anterior thalamic radiations, and cingulum were significantly different between the VPT-abnormal and the FT group. The VPT-normal group were found to have values between the VPT-abnormal and FT groups, but their values were not significantly different. This is in contrast to previous findings that have demonstrated differences between all three groups. This may indicate that while white matter is affected by preterm birth, greater damage to these fibers may be associated with later motor deficits.

A major finding of this study is the relationship between corpus callosum anatomy and the MABC-2 in VPT-born children with motor impairment. A study using comparable methods demonstrated a relationship between diffusion properties of the corpus callosum and MABC-2 scores in typically developing preschoolers (Grohs, Reynolds, Dewey, & Lebel, 2018). They concluded that a more a mature corpus callosum, as reflected by higher FA and lower radial diffusivity, may account for better motor skills in typically developing children. The lower FA and higher MD in the forceps major fibers found in the VPT-abnormal group in the current study may therefore reflect less mature white matter. The higher FA and lower MD in the FT and VPT-normal groups may indicate that these children have increased myelination and/or a decrease in extracellular space associated with a more typical developmental trajectory (Qiu, Mori, & Miller, 2015).

We also demonstrated lower brain stem volume and higher ventricle volumes across VPT groups compared to the FT group. This finding is in line with previous findings in older children who were born preterm (de Kieviet et al., 2012; Solsnes et al., 2016) and likely represents areas that are vulnerable to damage caused by early birth, rather than specific to movement abilities.

The current study does have limitations. This was a small convenience sample of low-risk VPT children without significant medical complications, thus limiting the generalizability of these results to the overall population of VPT-born children. The children also had a range of gestational ages and birthweights. Studies performed in higher risk populations (e.g., > 28 weeks gestation or < 1000 gm birthweight) may reveal more significant abnormalities.

We have shown that healthy preschool children born VPT who demonstrate motor deficits have neuroanatomical differences compared to their age-matched normal VPT-born peers. These neuroanatomical regions may be important markers for predicting longer term outcomes. It is possible that the motor impairments present at preschool age and the correlations between movement skills and measures of brain structures and integrity will persist as children grow. It is also possible that motor deficits may become more profound as older children are expected to complete become more complex motor tasks.

Early motor development has been demonstrated as an important predictor of future cognitive skills (Oudgenoeg-Paz, Mulder, Jongmans, van der Ham, & Van der Stigchel, 2017). Motor skill weaknesses may thus provide clues regarding which VPT children are at risk for future cognitive challenges. This study emphasizes the importance of longitudinal neurodevelopmental follow-up of VPT children since motor difficulties in VPT children without previously identified neurodevelopmental impairments may not become evident unless formally assessed in the preschool period. Further follow-up studies are needed to examine the relationship between motor skills at preschool age and scores on measures of movement, academic, and cognitive function in later childhood and adolescence. Early recognition of motor deficits may enable early intervention that can improve motor and academic outcomes in children born very preterm (Litt, Glymour, Hauser-Cram, Hehir, & McCormick, 2017).

## Compliance with Ethical Standards

### Funding

This study was funded by the *Eunice Kennedy Shriver* National Institute of Child Health and Human Development under Grant R01HD075865 (Akshoomoff) and in part by Grants R01HD061414 (Jernigan) and RC2DA029475 (Jernigan).

### Ethical Approval

All procedures performed in studies involving human participants were in accordance with the ethical standards of the institutional (University of California, San Diego) committee and with the 1964 Helsinki declaration and its later amendments or comparable ethical standards.

